# Inflammasome activation dictates the efficacy of antimycobacterial activity of frontline TB drugs

**DOI:** 10.1101/2025.04.02.646750

**Authors:** Anjali Singh, Kanika Bisht, Nidhi Yadav, Ranjan Nanda, Vivek Rao

## Abstract

Recent developments in TB treatment have identified an enormous potential of host targeting entities (host directed therapies-HDT) in achieving better and faster control of infection. We have previously demonstrated the synergistic effect of sertraline with frontline TB drugs in clearing infection in murine tissues. We surmised that understanding the mechanism of sertraline mediated enhancement of antibiotic efficacy would help identify novel pathways of pathogen control. Here, we have used sertraline as a probe to identify host signaling pathways that are important for bacterial control. We identify a role for sertraline mediated modulation of macrophage mitochondrial physiology in this process. We show that the resultant ROS generated in cells acts as the secondary signal for host cell inflammasome activation catalyzing the greater release of IL1β and significant efflux of K^+^ from the SRT treated macrophages. We thus highlight an important relationship between mitochondrial physiology in regulating the activation of the inflammasome to achieve better control of Mtb by infected macrophages.

**Graphical abstract:** 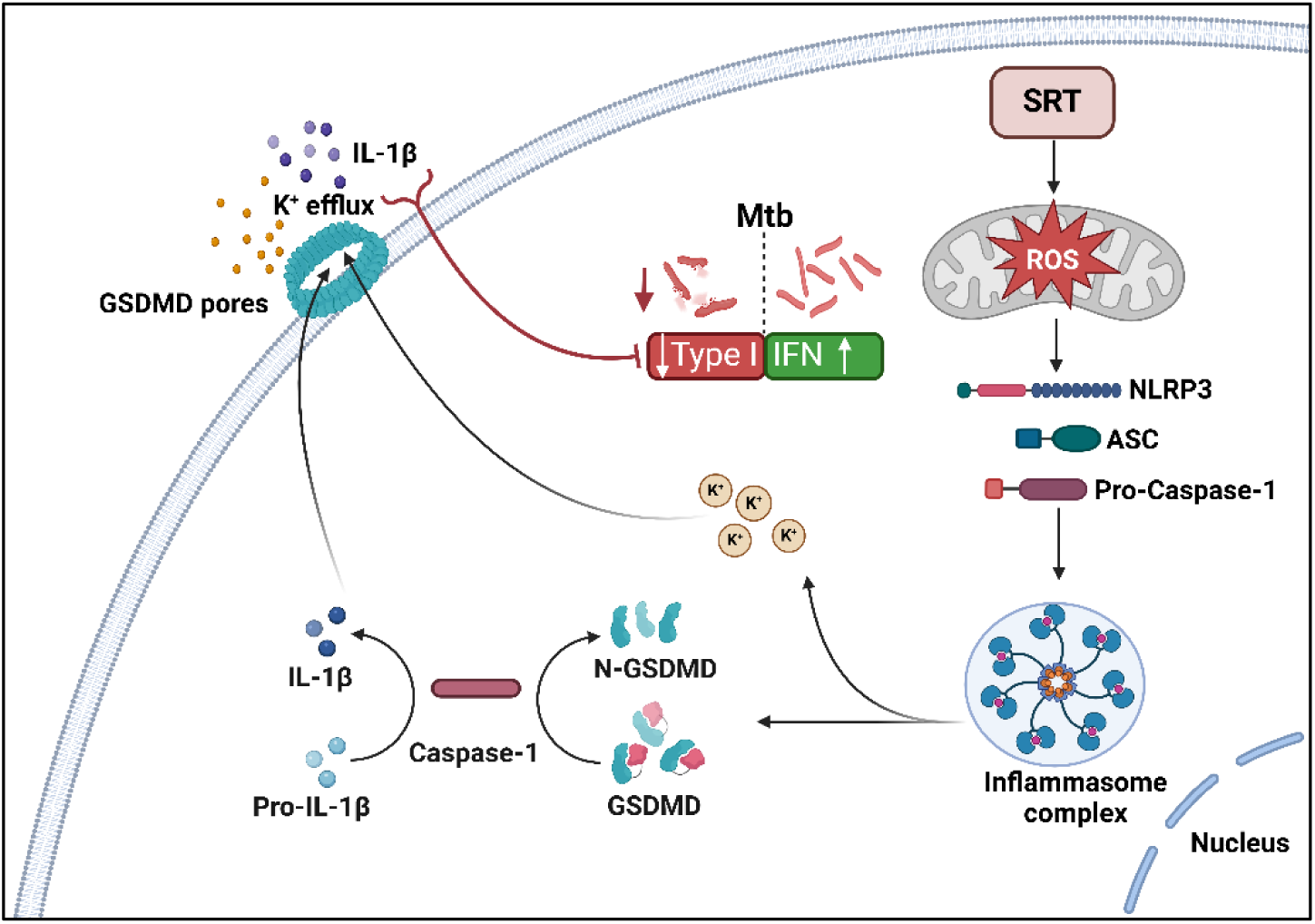

## Introduction

The current trends towards TB control have centered on harnessing the host responses for bacterial control as an effective stratagem to circumvent drug-associated development of resistance in the population [1, 2]. Several modalities have been tested as host-directed therapies for improved bacterial control, accelerated bacterial clearance, and symptomatic relief [3, 4]. We have previously demonstrated that the addition of an FDA-approved antidepressant drug-sertraline (SRT) significantly enhanced Mtb growth control in cellular and animal models of infection conferring higher resistance to tissue damage, better host survival, and faster bacterial clearance from the tissues [5]. SRT, in addition to its property of downregulating the type I IFN responses, has previously been documented to influence multiple cellular and immune mechanisms such as induction of cytokines like IFN-γ and IL-10, modulating autophagy and apoptosis, altering natural killer (NK) cell stimulation, and the inhibition of cellular translation machinery [6–9]. These pleiotropic effects of SRT extend beyond its primary function as a neurotransmitter and suggest a broader impact on host immune responses.

We have previously demonstrated host cell inflammasome activation by SRT as an important contributor to its antibacterial properties. The host cell inflammasome is now recognized as a key regulator of macrophage innate defense against infections. The significantly higher mortality and disease severity in IL-1β- or IL-1 receptor-deficient mice supports the necessity of inflammasome activation dependent production of IL-1β in control of Mtb infection [10–12]. We further validate the relevance of the inflammasome activation by systematically inhibiting the individual components of the activation pathway resulting in the activation of the macrophage Gasdermin D (GSDMD) by SRT. Although, the importance of gasdermins - D, E, C in the release of these mature cytokines and in the control of infections is well established, its relevance in Mtb clearance is unclear [13, 14]. Here, we conclusively establish the importance of GSDMD in antibiotic-mediated control of Mtb in both cellular and *in vivo* models of infection and deposit the consequent efflux of K^+^ ions as a key factor in hampering the pathogen-beneficial type I IFN response in Mtb infections.

The NLRP3 inflammasome is known to be activated by a two-stage mechanism, the first trigger (priming) is linked to NF-κB activation resulting in NLRP3, pro-IL1β/ IL-18 expression and the ensuing activation of NLRP3 for the formation of the mature inflammasome complex as the second step resulting in the maturation and release of active IL1β and IL18 via gasdermin D (GSDMD) pore [15–18]. With infection acting as the primary trigger, we uncover the significance of mitochondrial perturbation in SRT-treated cells as the secondary triggers for inflammasome activation thus highlighting a novel and important role for cellular physiology in the control of infections by antibiotics.

## Methods

### Reagents

THP1 Dual Monocytes and gasdermin D knockout monocytes (thp-kogsdmdz) were obtained from InvivoGen (Toulouse, France). The integrity of the cells was tested by STR profiling with routine mycoplasma testing. The cells were grown in HiglutaXL RPMI-1640 and 10% Fetal Bovine Serum (HiMedia laboratories, Mumbai, India), and differentiated to macrophages with PMA (Phorbol 12-Mysristate 13-acetate-P8139, Sigma Aldrich, USA) for 24h. The following reagents were procured from Sigma Aldrich, USA: Isoniazid (I3377), Pyrazinamide carboxamide (P7136), Ethambutol dihydrochloride (E4630) and Sertraline hydrochloride (S6319). Rifampicin (CMS1889, HIMEDIA laboratories, Mumbai, India) and commercially available SRT (Daxid, Pfizer Ltd, India) were used for mouse studies. NLRP3 inhibitor MCC950 (10µM) (Cayman Chemical Company), Caspase-1 inhibitor VX-765 (EMD Millipore Corp), Caspase-4 inhibitor IHC-2 (Sigma-Aldrich, USA), GSDMD inhibitor disulfiram (DSF; Sigma-Aldrich, USA), dimethyl fumarate (DMF; Sigma-Aldrich, USA) and for potassium efflux inhibition, KCl (50mM) (Sigma-Aldrich, USA) was used. For activation of inflammasomes, cells were treated with lipopolysaccharide (LPS; InvivoGen, France) and Nigericin (Cayman Chemical Company). Rotenone (Sigma-Aldrich, USA), CCCP (carbonyl cyanide m-chlorophenylhydrazone) (Sigma-Aldrich, USA) and MitoSox dye (5 µM) was purchased from (Thermo Fisher Scientific). For labelling mitochondria, mitotracker deep red at (500nM) dye was utilised (Thermo Fisher Scientific).

### Culture of bacteria and cell lines

THP-1 monocytes were cultured in RPMI 1640 supplemented with 2 mM L-glutamine, 25 mM HEPES, 50 µM sodium pyruvate, and 10% fetal bovine serum at 37^0^C in a 5% CO₂ humidified incubator. PMA (Phorbol 12-Mysristate 13-acetate-P8139, Sigma Aldrich, USA) was used for differentiation into the macrophages for 24h [19]. For isolation of human monocyte derived macrophages (MDM), PBMCs isolated from healthy volunteers were grown in RPMI containing 50ng/mL of granulocyte-macrophage colony-stimulating factor (GM-CSF) (Invitrogen) for 7 days in accordance with Institutional human ethics committee approval (Ref no: CSIR-IGIB/IHEC/2017–18). The macrophages were then removed from the plates and used for infection studies.

For infections, Mtb Erdman was cultured in Middlebrook 7H9 broth (BD, Difco), supplemented with 0.5% glycerol, 4% Albumin-Dextrose-Saline (ADS), and PBST (0.05% Tween 80) at 37^0^C. Bacterial numbers were calculated in the single cell suspensions (SCS) and used to infect macrophages at MOI 5 in the presence / absence of the inhibitors for 6h. Following this, cells were either left untreated or treated with antibiotics (H-20ng/ml and R-100ng/ml) with and without SRT-20µM and the bacterial numbers were estimated by plating serial dilutions of cell lysates on Middlebrook 7H10 agar containing 0.5% glycerol and 10% Oleic acid-Albumin-Dextrose-Catalase (OADC) supplement (HiMedia Laboratories, India) and incubation at 37^0^C for 3-4 weeks.

### Treatment of the Inflammasome modulators

Macrophages were pre-treated for 30 minutes with inhibitors, MCC950 (10µM), VX-765 (20µM), IHC-2 (20µM), DSF (10µM), DMF (50µM), and KCl (50mM) followed by infection with Mtb. For the activation of inflammasomes, cells were treated with 100ng/ml LPS for 3h and with nigericin for 45 min prior to infection with Mtb.

### Analysis of macrophage responses

The expression of cytokines in the cell supernatants was analysed by specific ELISA according to the manufacturer’s recommendations (IL1β, TNFα, BD OptEIA - BD biosciences), IFNβ (R&D systems).

The levels of IFNβ were estimated in the THP1 dual cells by quantitating the luminescence of the cell supernatants following the addition of the substrate-QUANTI-Luc (Invivogen, France), as per the manufacturer’s recommendations. GSDMD expression was detected by immunoblotting with a monoclonal antibody HPA04487 (*Prestige Antibodies*® Sigma-Aldrich, USA) and fluorescently labelled secondary antibodies and visualized with a LI-COR Odyssey imaging system.

For estimation of mitochondrial ROS, SRT-treated cells were incubated with 5 μM of the specific dye**-**MitoSOX for 30 min and imaged in the Invitrogen™ EVOS™ M5000 Imaging System. Cells treated with (rotenone (10 μM) was used as positive controls for mitochondrial ROS, For analysing the mitochondrial potential, THP1 cells were treated with SRT for different time intervals, stained with 1µM tetramethylrhodamine (TMRE,) (Thermo Fisher Scientific) at 37 °C for 30 minutes and analyzed by FACS (BD Accuri™ - BD Biosciences). CCCP at 10 μM (Sigma-Aldrich, USA) was used as a control for the study.

### Mtb infection in mice model of infection

All animal infections were conducted in a dedicated ABSL-3 facility as per the accepted recommendations of the IAEC (IGIB/IAEC/10/Nov/2023/05). BALB/c mice (aged 6-8 weeks) were infected with Mtb using an inhalation exposure system (Glas-Col, USA) for aerosol delivery of ∼ 500 cfu per animal. Treatment with antibiotics and SRT was initiated after 4 weeks of infection and provided *ad libitum* in the drinking water containing 1% sucrose twice a week. Mice were administered the TB drugs (H-100 mg/kg, R-40 mg/kg, Z-150 mg/kg, E-100 mg/kg), with or without SRT (10 mg/kg) and in the presence or absence of DSF (300 mg/kg). At specific time points, animals were euthanized and the tissues were used for estimation of the bacterial numbers and histological examination. The formalin-fixed left caudal lung lobes were used for gross examination by a Zeiss (Semi 2000®C) bright field microscope and histological evaluation by H&E staining of sections in an Olympus microscope.

### Analysis of cellular potassium levels

Intracellular or excreted potassium levels were estimated in SRT-treated THP1 cells by two methods-Mass spectrometry and staining with a fluorescent potassium-specific dye-ION potassium green-4 acetoxymethyl ester (IPG-4 AM; Cayman Chemical Company). For the latter method, cells were treated with the dye in 0.5% (w/v) pluronic acid F-127 (Sigma-Aldrich, USA) for 1h at 37 °C in a CO_2_ incubator in the dark and then imaged using the Invitrogen™ EVOS™ M5000 Imaging System. For mass spectrometric estimation, THP1 cells at a density of 60,000 cells were treated with SRT in isosmotic buffer K (130mM NaCl, 7mM Na_2_HPO_4_, 3mM NaH_2_HPO_4_. 5mM KCl, pH 7.4) for 3h. After washing cells with ICP grade water twice, the pellet was resuspended in 150µl of 70% HNO_3_ and 50 µl of 30% H_2_O_2_ and incubated in the dark for 10 min. Following digestion with HNO_3_ and H_2_O_2_ (ramp = 250ω for 10 min, Hold = 250ω for 5 min and cool = 55 °C), the samples were filtered and analyzed in the ICP-MS machine (iCAPTM TQ ICP-MS, Thermo Scientific, USA) according to the manufacturer’s protocols. A standard curve was used to estimate the concentration of K^+^ ion in the samples. Cells incubated in buffer W without KCl and with 50mM KCl were used as a positive and negative control for K^+^ efflux respectively.

### Statistical tests and significance

All statistical analyses were conducted using GraphPad Prism software. An unpaired t-test or multiple t-tests were applied to data pooled from two to three independent experiments, each performed in triplicate. For parametric data analysis, the student t-test with Welch’s correction method or ordinary one-way ANOVA for multiple comparisons was used to determine the significance of the t-tests. For non-parametric data analysis statistical analysis was performed using a two-tailed Mann-Whitney test or Kruskal-Wallis test for multiple comparisons.

## Results

### Inflammasome activation is critical for enhanced antibiotic efficacy in Mtb infection

The importance of host cell inflammasome activation in controlling infections, specifically in the context of Mtb control is not clearly understood. While activation of this signaling pathway by Mtb infection is undebatable, studies have revealed host beneficial as well as a pro-pathogenic innate phenomenon in macrophages [20–22]. Our previous study suggested an important role of the host cell inflammasome in augmented control of Mtb by a combination of sertraline with frontline TB drugs [23].

To investigate whether inflammasomes determine antibiotic efficacy, we first tested the effect of heterologous activation of inflammasomes by LPS and nigericin on bacterial control. The addition of LPS and nigericin (L+N) prior to infection alone was sufficient to restrict Mtb growth early (day 1) by 3-folds that amplified to > 10 folds difference in the L+N group as compared to the untreated groups by day 3 (Figure 1A). HR also was able to significantly decrease bacterial burdens in macrophages by 1.5 folds by day 1, a combination with LPS and nigericin enhanced this effect to 4 folds. Even after 3 days of infection, the combination of LN and HR enhanced growth control by 2.7 folds in comparison to HR alone corroborating a synergistic effect of inflammasome activation on antibiotic efficacy.

**Figure 1:**
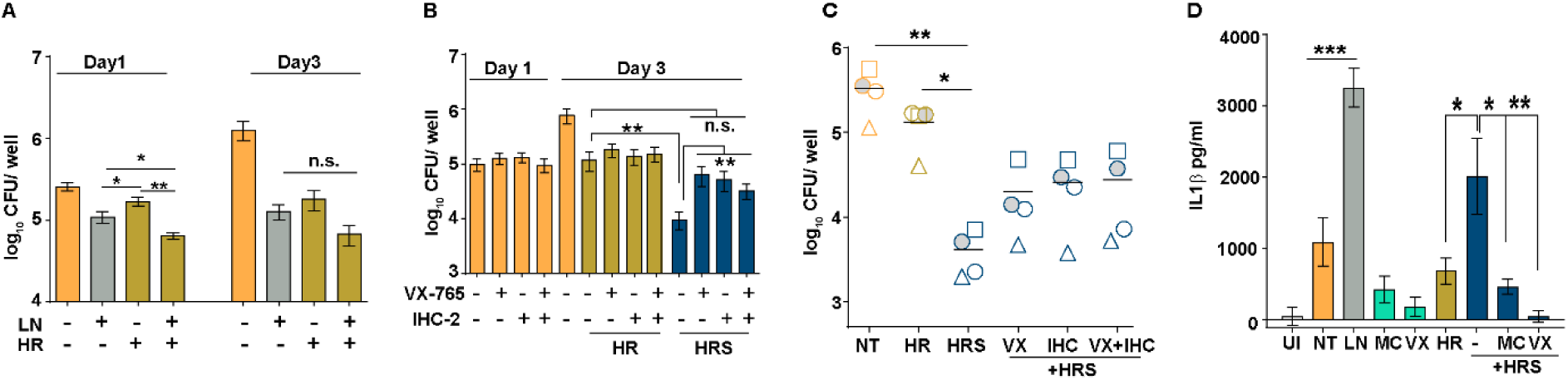
Sertraline-mediated enhancement of antibiotic efficacy is dependent on inflammasome activation. A) The effect of LPS and nigericin (LN) on bacterial growth in THP1 macrophages was evaluated and compared to the effect on the bacterial control by frontline TB drugs (HR). Bacterial numbers at day1 and day3 post treatment were enumerated by CFU plating and are represented as mean CFU ± SEM of triplicate assays from 3 independent experiments (N=3). B, C) Bacterial growth in THP1 (B) or human monocyte-derived macrophages (C) that were infected with Mtb and either left untreated (NT) or treated with HR, HR+ sertraline (HRS) alone or in combination with caspase 1 (VX765)/ caspase 4 specific (IHC-2) inhibitors. Bacterial numbers at day 1 and day 3 post-treatment (B) and day 5 post-treatment (C) were enumerated by CFU plating and are represented as mean CFU ± SEM of triplicate assays from 3 independent experiments (N=3). Each symbol in panel C represents a sample from one healthy individual. D) The levels of secreted IL1β in culture supernatants of Mtb infected THP1 macrophages left untreated (NT) or treated with HR, HR+ sertraline (HRS) alone or in combination with caspase 1 (VX765-VX)/ NLRP3 (MCC950-MCC) inhibitors. The level of IL1β was measured after 24h of infection by ELISA and is represented as mean concentrations (pg/ml) ± SEM of triplicate assays from N=3 experiments.

We have previously shown that the effect of SRT in augmenting Mtb control can be reversed by the inhibition of the NLRP3 inflammasome by MCC950 [23]. To investigate the complete cascade of inflammasome activation, we inhibited inflammasome-associated caspase activation and tested the effect of sertraline on bacterial control. While SRT alone enhanced HR-mediated control of Mtb in macrophages by 6.5 folds on day 3, the addition of VX765 or IHC-2, inhibitor of caspase1 and caspase 4, respectively, despite not altering the initial uptake, completely reversed this increment with bacterial numbers reaching levels comparable to HR treated macrophages (Figure 1B). This effect of inhibition of caspase on sertraline effectivity was also observed in primary human monocyte-derived macrophages wherein both IHC-2 and VX765 reversed the heightened bacterial control observed in HRS treatment in all the individual samples implying the importance of caspase/inflammasome activation (Figure 1C). Inflammasome activation results in the secretion of mature IL1β and IL18 from activated macrophages [24]. Infection with Mtb induced significant levels of IL1β in macrophages (∼1000pg/ml). While treatment with HR did not alter secretion, the addition of SRT induced ∼2-fold higher levels of IL1β secretion in the macrophages (Figure 1D). In line with previous reports, the IL1β levels in macrophages treated with LPS and nigericin were significantly ∼3 folds) higher than infection alone. Addition of the inflammasome inhibitors-MCC950 or VX765, significantly reduced IL-1β secretion.

### Sertraline-induced gasdermin D activation is responsible for higher IL1β secretion in macrophages

Inflammasome initiation leads to the cleavage of the membrane pore-forming molecule-gasdermin D (GSDMD) and the release of pro-inflammatory cytokines. To investigate the role of GSDMD in antibiotic potentiation, macrophages were treated with two distinct GSDMD inhibitors: dimethyl fumarate (DMF), an inhibitor of the cleavage of the inactive GSDMD into its pore-forming active form or disulfiram (DSF), a potent inhibitor of GSDMD pore formation along with HRS. Despite a negligible impact on the initial uptake of Mtb (d0), addition of DSF or DMF reduced the effect of sertraline in augmenting HR-dependent bacterial control (Figure 2A). In contrast to a 4-5-fold decrease in bacterial numbers by day3 with HR, HRS treated macrophages further reduced these numbers by 6-8 folds. Macrophages treated with DSF along with HRS supported increased bacterial growth by 3-4 folds while the addition of DMF to HRS increased the bacterial numbers to almost the levels of HR alone (negating the effect of SRT completely), validating the importance of this host signaling component in enhanced antibiotic efficacy. This was also evident as a loss of the effect of SRT on HR activity in primary human monocyte-derived macrophages from healthy individuals with the addition of either DSF or DMF (Figure 2B). A significant increase in gasdermin D activation was observed by SRT treatment (Figure 2C). Treatment of macrophages with either SRT or LPS+ Nigericin (LN), a potent activator of inflammasome, enhanced the formation of the 31kDa N-terminal active form of GSDMD that was abrogated in cells treated with DMF (Figure 2C). Given the important role of GSDMD in the release of mature cytokines from activated macrophages [13] we analyzed IL1β levels in the supernatants of infected macrophages treated with HRS and DMF or DSF. As expected, a significant 2.5-fold increase was observed in IL1β secretion by infected macrophages that were treated with HRS as compared to the untreated and HR-treated cells; this increment was completely abolished by the addition of the GSDMD activity inhibitors-DSF/ DMF either alone or in combination with HRS (Figure 2D). Interestingly, treatment of human MDMs with SRT induced a similar phenotype of cell swelling with the formation of large fluid-filled vacuolar structures in macrophages as described earlier [25, 26] (Figure S1A). This effect was completely reversed by the addition ofDSF, implicating the action of SRT in activating host cell inflammasomes via the GSDMD signaling pathway.

**Figure 2:**
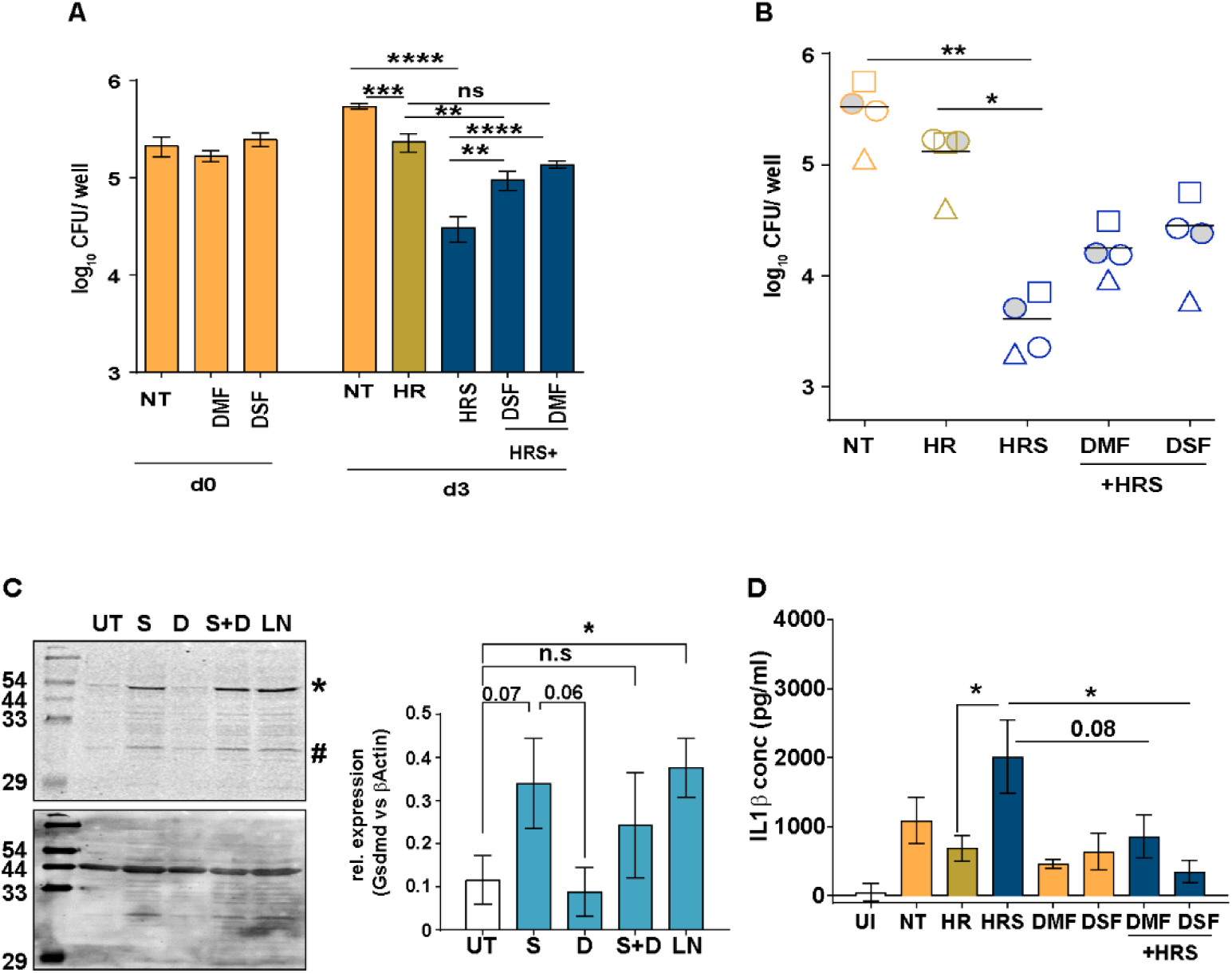
Sertraline activates gasdermin D in treated macrophages. A, B) Bacterial growth in THP1 (A) or monocyte-derived macrophages (B) that were infected with Mtb and either left untreated (NT) or treated with HR, HR+ sertraline (HRS) alone or in combination with gasdermin D activation inhibitors-dimethyl fumarate (DMF)/ disulfiram (DSF). Bacterial numbers at day0 and day3 post treatment (A) and at day 5 (B) were enumerated by CFU plating and are represented as mean CFU ± SEM of triplicate assays from 3 independent experiments (N=3). Each symbol in panel B represents a sample from one healthy individual. C) Expression levels of mature gasdermin D in crude lysates from infected macrophages treated with LPS+ nigericin (LN) or DMF at the indicated concentrations. The premature form (*) and the mature N-terminal form (#) were analyzed by immunoblotting with gasdermin D-specific antibody. Quantitation of the mature form of gasdermin D by densitometric analysis, normalized to β-actin, is shown in panel C.

### GSDMD activation is critical for increased antibiotic efficacy

Previous reports have demonstrated the essentiality of GSDMD in controlling infection in macrophages [27–29]. In sync with these observations, the importance of GSDMD was assessed in the context of Mtb infection. Mtb infected GSDMD^-/-^ macrophages were impaired in their ability to restrict bacterial growth as efficiently as wildtype (Wt) macrophages harboring 4-5-fold higher bacteria by day 3 of infection (Figure 3A). Interestingly, loss of GSDMD did not alter the ability of HR to control bacteria, the augmentation with SRT was significantly altered in these two cell types (Figure 3B). Cells deficient for GSDMD did not display increased bacterial control with HRS treatment as opposed to the Wt cells that showed more than 10-fold reduction in bacterial numbers when compared to the HR treated cells.

**Figure 3:**
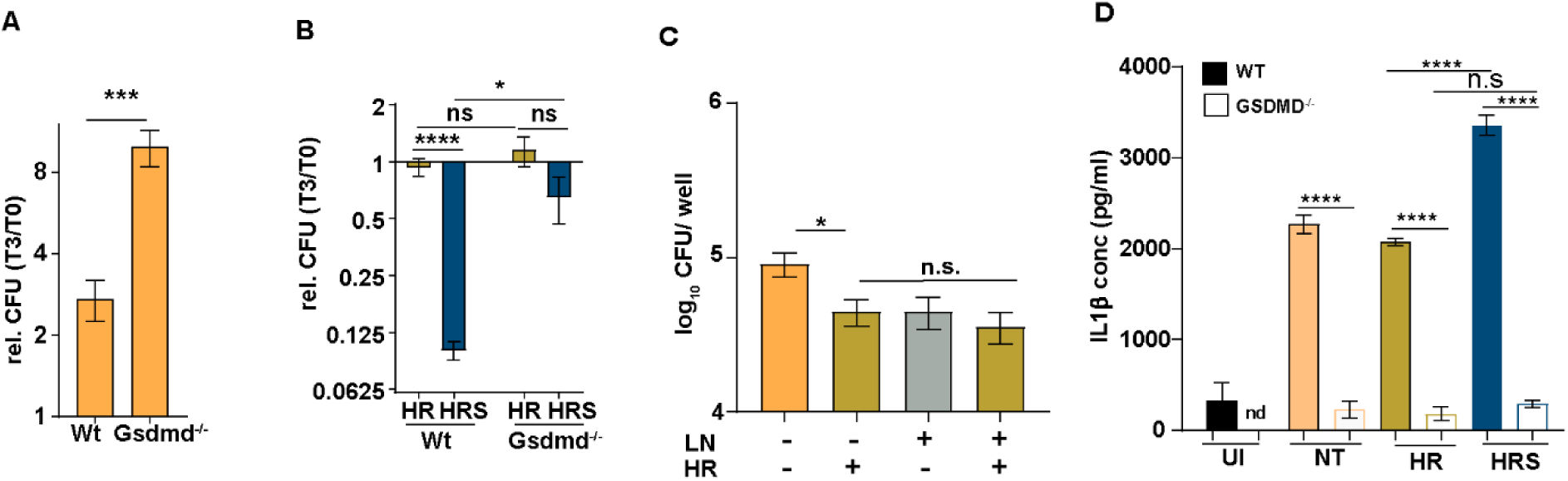
Gasdermin D activation is critical for Mtb control by infected macrophages. A, B, C) Bacterial growth in Wt or GSDMD deficient THP1 macrophages that were infected with Mtb and either left untreated (NT-A) or treated with HR, HRS (B), or in macrophages pretreated with the inflammasome activator LN (C). Bacterial numbers were enumerated by CFU plating and are represented as mean CFU ± SEM of triplicate assays from 3 independent experiments (N=3). D) The levels of IL1β secreted from Wt or GSDMD deficient THP1 macrophages infected with Mtb and left untreated or treated with HR, HRS. The level of IL1β was measured after 24h of infection by ELISA and is represented as mean concentrations (pg/ml) ± SEM of triplicate assays from 2 independent experiments (N=2).

The increased control observed with the activation of inflammasome with LPS and nigericin observed in HR treated Wt macrophages (Figure 1A) was lost in the GSDMD^-/-^ macrophages (Figure 3C) emphasizing the importance of this axis in bacterial control. This was further evident in the GSDMD^-/-^ macrophages that elaborated significantly lower levels of IL1β in response to infection at 18h of treatment (Figure 3D). Moreover, the observed difference in cytokine secretion between HR and HRS in the Wt cells was completely lost in the GSDMD^-/-^ cells further corroborating the involvement of inflammasome by SRT to enhance bacterial control in macrophages.

### GSDMD is critical for *in vivo* control of Mtb

To test the importance of GSDMD *in vivo*, Balb/c mice were infected with Mtb and treated with antibiotics along with the gasdermin inhibitor – disulfiram. The addition of disulfiram did not significantly alter the Mtb growth kinetics alone or in combination with frontline TB drugs (HRZE) by 8w of infection (Figure 4A). However, the effect of DSF treatment was pronounced in HRZES-treated animals. While the addition of sertraline to HRZE enhanced the control of Mtb in the lungs of animals, the inclusion of DSF resulted in a complete reversal of this added advantage leading to bacterial numbers similar to antibiotics alone. Moreover, this inhibitory effect of DSF was also evident in tissue damage in animal lungs (Figure 4B). As expected, H&E-stained histopathological analysis of the lungs from HRZES-treated animals showed fewer microscopic lesions compared to untreated or HRZE-treated animals. However, sections from animals treated with HRZES along with DSF showed significantly higher lesions, similar to those in the HRZE-treated or untreated groups (Figure 4B)

**Figure 4:**
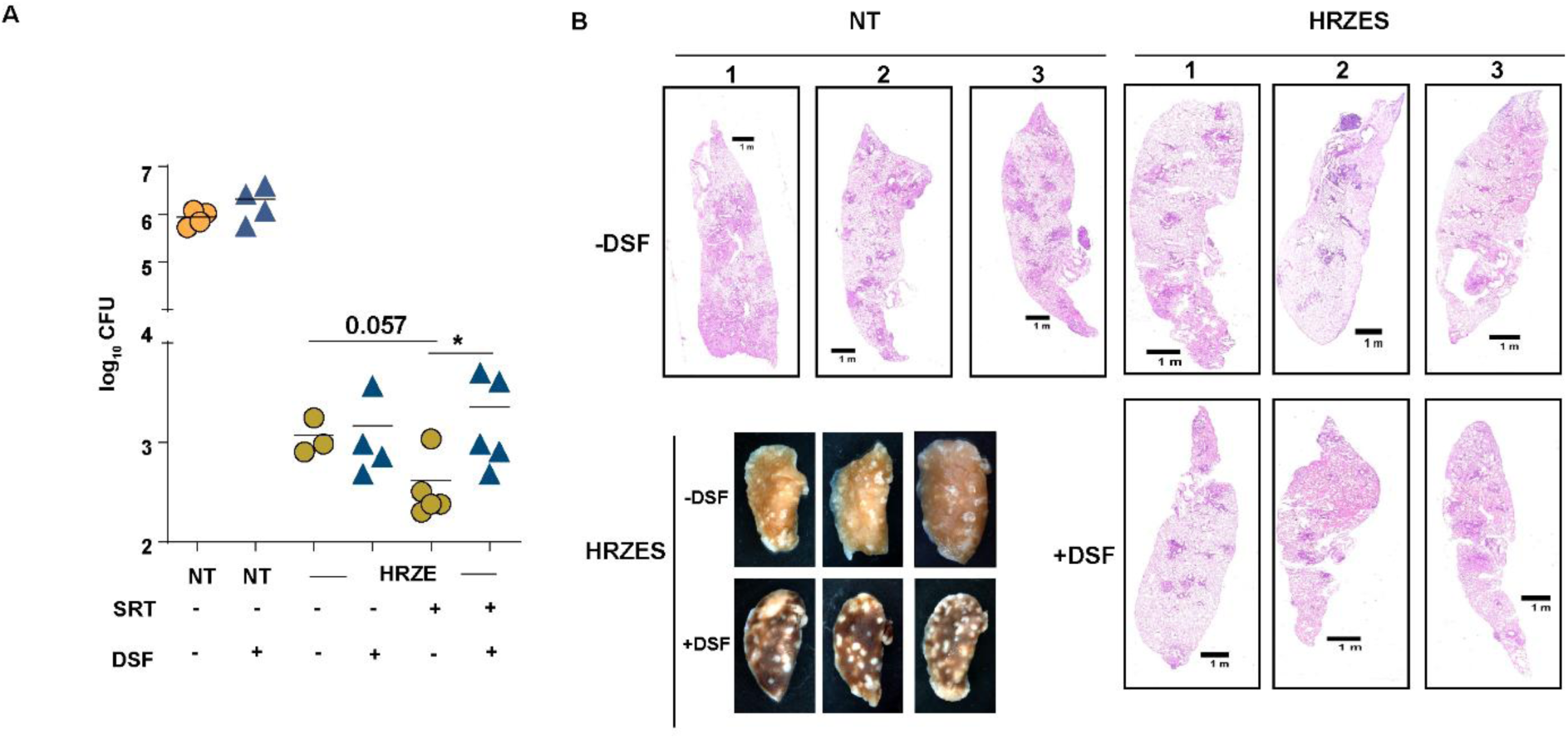
*In vivo* Gasdermin D activation dictates Mtb control. A) Bacterial growth in lungs of Balb/c mice that were infected with 500 cfu of Mtb by aerosol and either left untreated (NT) or treated with HRZE, HRZE+ sertraline (HRZES) alone or in combination with disulfiram (D, HRZED, HRZESD). Bacterial numbers at 4w were enumerated by CFU plating and are represented as mean CFU ± SEM of lungs from 4 or 5 mice per group. B) Macroscopic images of lung lobes from HRZES ± DSF treated animals and histological examination of lung sections stained with H&E from NT – DSF and HRZES ± DSF treated animals.

### Inflammasome-mediated potassium efflux regulates Mtb-induced type I IFN responses and is critical for enhanced bacterial control

One of the consequences of gasdermin D activation is the release of intracellular potassium ions [30]. To understand the relevance of this efflux, the levels of intracellular/ secreted potassium were estimated in THP1 macrophages following SRT treatment. Treatment with nigericin, a potent inflammasome activator induced significant levels of potassium release from the cells showing 4-5 folds lesser potassium compared to untreated cells with concomitant excess in the supernatants of these cells (Figure 5A). Cells incubated with a potassium ion-depleted buffer (W) also facilitated K+ ion efflux with levels decreased by 6-7 folds, serving as a control. Treatment with SRT also induced significant efflux as evidenced by the lower intracellular and higher extracellular concentrations of potassium in macrophages. While the addition of KCl to the medium did not alter the intracellular levels in untreated cells, the efflux was completely abolished in cells with KCl. In order to understand the kinetic flux of K^+^ in macrophages, cells were stained with a specific dye IPG-4 AM and imaged after treatment. The ability of SRT to induce K^+^ efflux was evident by 6h that gradually increased by 48h of treatment evidenced by the significantly lower fluorescence in comparison to untreated cells. A similar phenotype was also observed for nigericin with significantly lower fluorescence in these cells between 6 and 48h of treatment. To determine the requirement of GSDMD pore formation for K⁺ ion efflux, cells were treated with the GSDMD inhibitor DMF; while DMF alone induced greater intracellular K^+^ concentrations at all time points, addition of SRT reversed this efflux to control levels suggesting that SRT induced gasdermin D activation resulted in the release of K^+^ ions (Figure 5B). Inhibition of SRT-mediated K^+^ efflux by the addition of KCl in the medium completely offset the extended bacterial control imparted by the addition of SRT in the HRS-treated macrophages, despite any influence on the bacterial control in HR-treated or untreated cells (Figure 5C). In line with these observations, macrophages lacking GSDMD did not extrude K^+^ ions following SRT treatment at any time of the experiment (Figure 5D) validating the importance of GSDMD in inducing K^+^ efflux from the macrophages.

**Figure 5:**
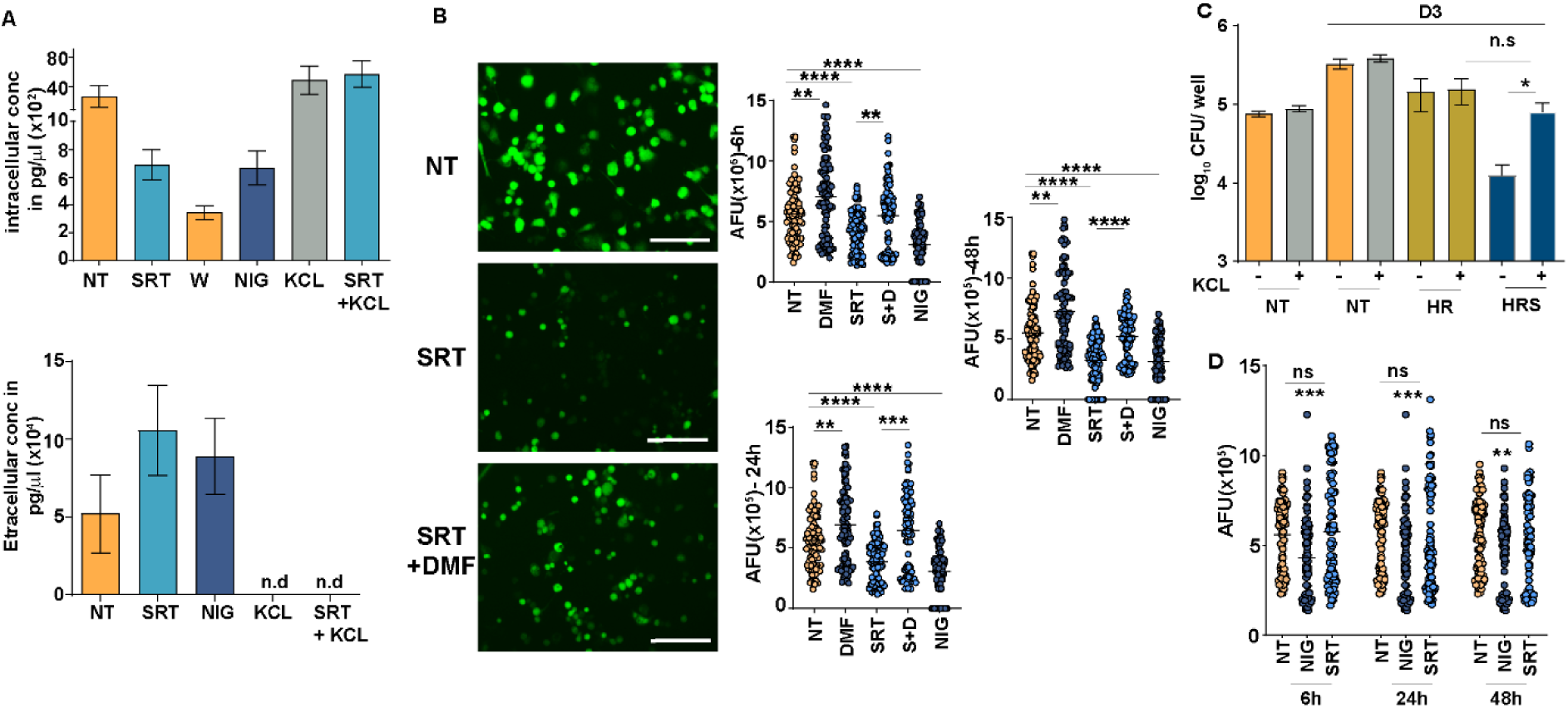
Sertraline-induced potassium efflux is critical for enhancing antimycobacterial control. A) The levels of potassium in cells left untreated (NT) or treated with sertraline (SRT) with and without KCL or nigericin (NIG) were estimated in macrophages and in the cell supernatants by ICP-MS. Values are mean ± SEM of 3 independent experiments. B) Estimation of potassium in macrophages left untreated or treated with SRT alone or with DMF as depicted by dye binding. Representative images of macrophages stained with IPG-4AM are shown while the kinetic change in potassium concentration at 6, 24, and 48h post-treatment is graphically represented. The fluorescent intensities (represented by dots) were calculated and are estimated as mean fluorescence ±SEM of 90 cells for 3 independent experiments expressed in arbitrary fluorescence units (AFU). C) The effect of KCl treatment on the ability of SRT to enhance bacteria growth control. Bacterial numbers at day 3 post-treatment were enumerated by CFU plating and are represented as mean CFU ± SEM of triplicate assays from 3 independent experiments (N=3). D) The extent of K^+^ ion efflux in GSDMD^-/-^ macrophages is depicted with the quantitative comparison of the temporal changes in these macrophages from the Wt cells are represented as mean CFU ± SEM of triplicate assays from 3 independent experiments (N=3).

### SRT-mediated K^+^ ion efflux regulates type I IFN response in Mtb infected macrophages

Given the significant release of K^+^ by SRT and our previous observations of SRT-mediated inhibition of type I IFN in macrophages, we investigated the role of inflammasome in this process. Mtb-infected macrophages were evaluated for the activation of the type I IFN response after treatment with inflammasome modulators. As expected, while Mtb infection activated type I IFN signaling in macrophages with minimal change on antibiotic treatment, the addition of nigericin alone or with HR significantly lowered these values by ∼6-10 folds by 24h of infection (Figure 6 A). In line with previous observations, SRT restricted the type I IFN signaling in macrophages by 3-4 folds, a property that was overturned by the addition of KCL alone or in combination of HRS to the medium (Figure 6A). In contrast to nigericin, inhibition of inflammasome by MCC950 or VX786 and GSDMD again significantly offset the restricted type I IFN response of HRS-treated cells (Figure 6B). While inhibition of inflammasome signaling was sufficient to restore the type I IFN levels of HRS-treated cells to untreated levels (Figure 6C).

**Figure 6:**
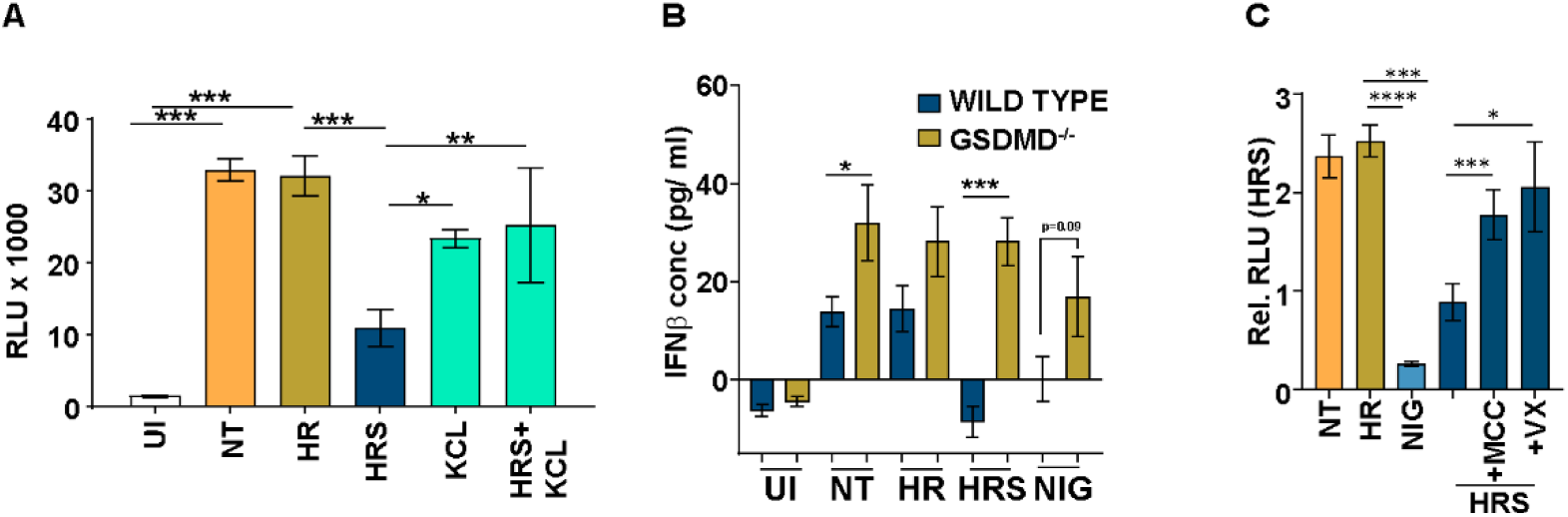
Inflammasome activation restricts Mtb induced type I IFN responses. A-C) The effect of inflammasome activation on the type I IFN response in control (UI) or Mtb infected THP1 dual macrophages, either untreated (NT) or treated with HR alone or in combination is depicted. A-Effect of K+ efflux inhibition with KCL, B-in Wt or GSDMD^-/-^ macrophages, C-in the presence of inflammasome activator-nigericin (NIG) or inhibition (MCC950 and VX765-VX). The extent of type I IFN regulated luminescence elicited at 24 h post infection (p.i.) is shown as relative units (RLU) ± SEM of triplicate assays from N=3 experiments.

### SRT induces significant alteration of mitochondria in macrophages

Inflammasome activation involves two stages with an initial NF-κB-dependent priming step, followed by a secondary activation signal that can stem from ER stress or mitochondrial dysfunction as one of the major triggers [31, 32]. To assess the role of ER stress as a potential trigger of inflammasome activation we evaluated the levels of XBP-1 splicing in cells treated with SRT or with tunicamycin (TM), a potent ER stress inducer. While tunicamycin induced near-complete splicing of XBP-1 as early as 1h with a gradual increase until the 24h after treatment, SRT failed to activate this process at any point of treatment (Figure 7A). Even Mtb-infected macrophages treated with HR or HRS did not show any evidence of ER stress as the spliced XBP-1 levels were undetectable in the extracts of these cells at any time point, suggesting that SRT does not induce ER stress. This was also evident in the complete lack of an effect on the alteration in bacterial control by antibiotics alone or in conjunction with SRT along with either the ER stress activator – tunicamycin or inhibitor – 4-phenylbutyric acid (PB) (Figure 7B). While HRS-treated cells showed decreased bacterial numbers than HR treatment, the addition of tunicamycin to HR or BP to HRS-treated cells did not alter the bacterial burdens even by day 3 suggesting that the SRT-mediated antibiotic efficacy is not due to ER stress-mediated inflammasome activation. In contrast, examination of macrophages treated with SRT revealed significantly elongated mitochondria in comparison to the untreated cells hinting at a possible role for an alteration of mitochondrial physiology as a trigger for inflammasome activation (Figure 7C). In fact, mitochondria in cells treated with SRT showed a significant increase in the extent of mitotracker deep red staining that is dependent on the membrane potential of the organelle indicating that SRT significantly alters the mitochondrial membrane (Figure 7D) [33]. This alteration of mitochondrial membrane potential was evident on TMRE staining of the cells, wherein treatment with a known oxidative phosphorylation uncoupler, (CCCP) results in mitochondrial membrane potential collapse with lower levels of staining (Figure S1B). The addition of SRT increases staining as early as 3h which is maintained by 24h of treatment indicating that SRT enhances mitochondrial membrane potential (ΔΨm), leading to hyperpolarization — a state often linked to increased electron transport chain (ETC) activity or mitochondrial stress. This sustained hyperpolarization would facilitate the generation of ROS leading to an altered mitochondrial metabolism and contributing to the downstream effects of inflammasome activation observed in SRT-treated cells.

**Figure 7:**
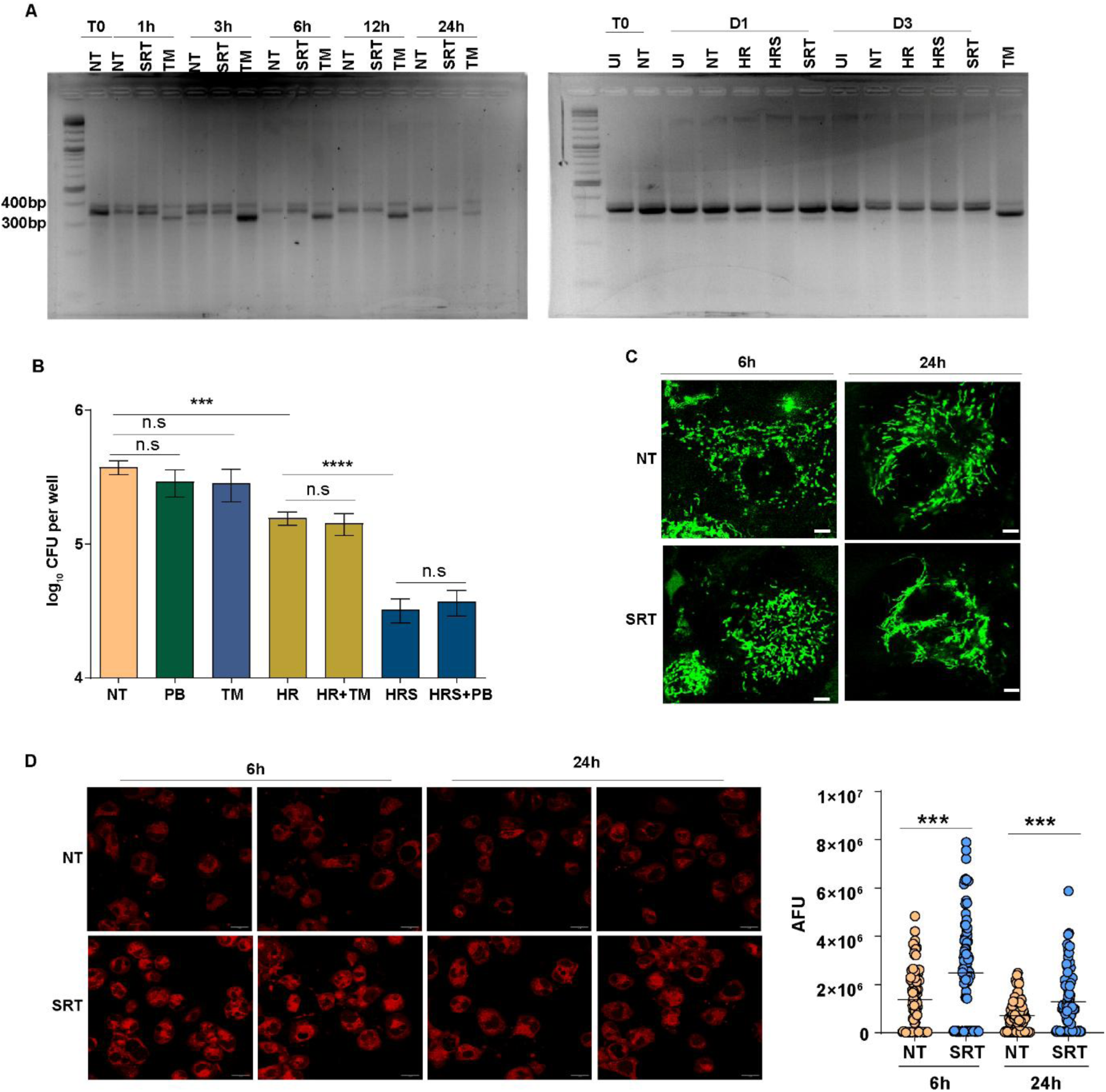
Sertraline induces significant alteration in the mitochondria of macrophages. A) The effect of sertraline on ER stress was analyzed by estimating the extent of XBP-1 splicing in uninfected (left panel) or the Mtb infected (right panel) macrophages and left untreated (NT) or treated with HR, SRT or HRS for the indicated time intervals. Cells treated with the potent ER stress inducer -tunicamycin (TM) were used as a positive control. B) The effect of modulating ER stress in cells by tunicamycin (TM) or 4-phenylbutyric acid (PB) on bacterial control by Mtb-infected macrophages. Bacterial numbers in THP1 macrophages that were infected with Mtb and either left untreated (NT-B) or treated with HR, or HR + sertraline (HRS) were enumerated at Day 0 and Day 3 post treatment and is represented as mean CFU ± SEM of triplicate assays from 3 independent experiments (N=3).C) Changes in mitochondrial morphology of THP1 macrophages treated with SRT for 6 and 24h were evaluated by staining with mitotracker green dye. Representative images of the cells (2) are depicted from 3 independent experiments (N=3). D) The extent of mitotracker deep red staining in untreated or SRT-treated macrophages was analyzed by microscopy. The mean arbitrary fluorescence units were calculated and are graphically represented as multiple cells/ fields from 3 independent experiments (N=3).

**Figure S1:**
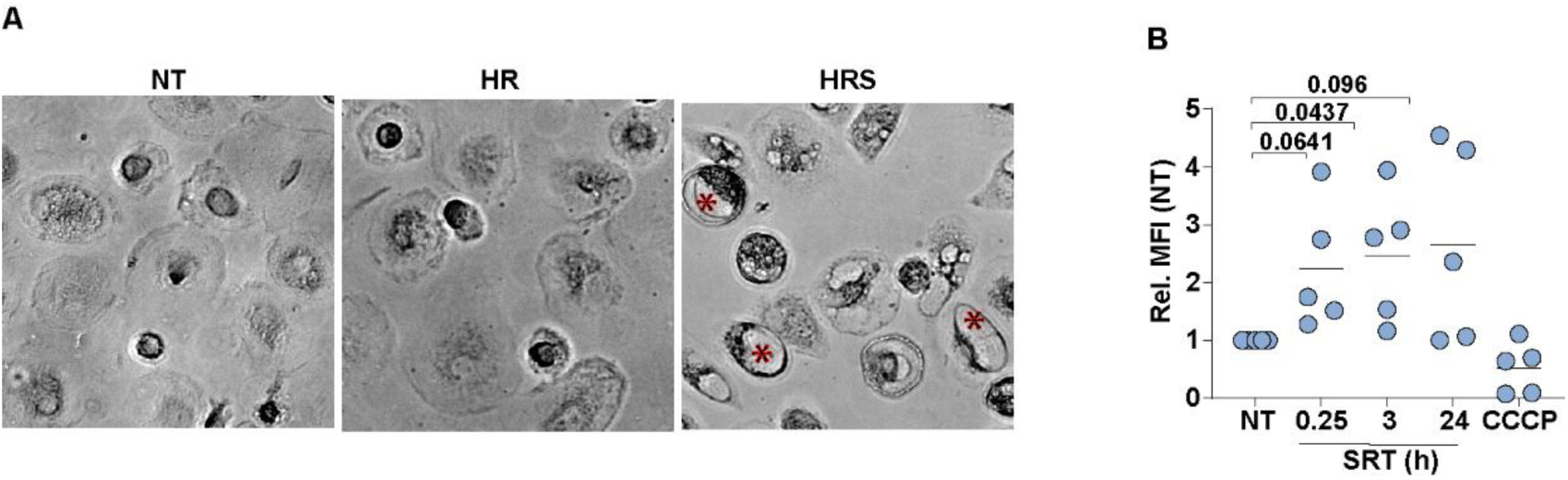
A) Effect of sertraline in human MDMs from healthy individuals. Mtb infected hMDMs were left untreated (NT) or treated with HR or HRS. A representative image of the cells after 5 days of infection is shown, the ballooning of cells is indicated by the red Asterix. B) The mitochondrial potential of THP1 cells treated with 20μM SRT for different time intervals as indicated was quantitated by FACS following staining with TMRE. The mean intensity values ± SEM of cells from 4 independent experiments is shown (N=4). Cells treated with the potent uncoupler CCCP were used as a control for effective loss of mitochondrial potential.

### Heightened mitochondrial ROS level in macrophages is critical for inflammasome activation in SRT treated cells

Normal mitochondrial function is crucial for cellular homeostasis. Disruptions in mitochondrial physiology can lead to significant alterations in overall cellular metabolism and response kinetics, particularly in innate immune cells like macrophages [34]. Several studies have highlighted the importance of mitochondrial ROS exacerbation in regulating multiple innate functions of macrophages, including inflammasome activation [34, 35]. With a significant alteration in mitochondrial architecture and potential by SRT, we analyzed the levels of mito-ROS and its role in the activation of the inflammasome. While the levels of MitoSox staining, a ROS-specific probe for mitochondria, in untreated cells were low, SRT treatment significantly enhanced the staining (Figure 8A). In fact, macrophages show significantly increased ROS as early as 6h that was maintained even after 48h of treatment with SRT at levels that are comparable to the potent mito-ROS inducer-rotenone. Inhibiting this ROS with a specific inhibitor-Mitoquinone (MQ),), a mitochondria specific ROS scavenger, reversed the enhanced bacterial control observed in the HRS treated cells without altering the ability of HR treated or the cells left untreated suggesting that mitochondrial ROS was important for the enhanced bacterial control mediated by SRT (Figure 8B). Furthermore, addition of menaquinone to macrophages treated with HRS significantly reversed the HRS-induced increase in IL-1β release, restoring these levels to those observed in untreated or HR-treated cells (Figure 8C) highlighting the pivotal role for mitochondrial perturbation in inflammasome activation.

**Figure 8:**
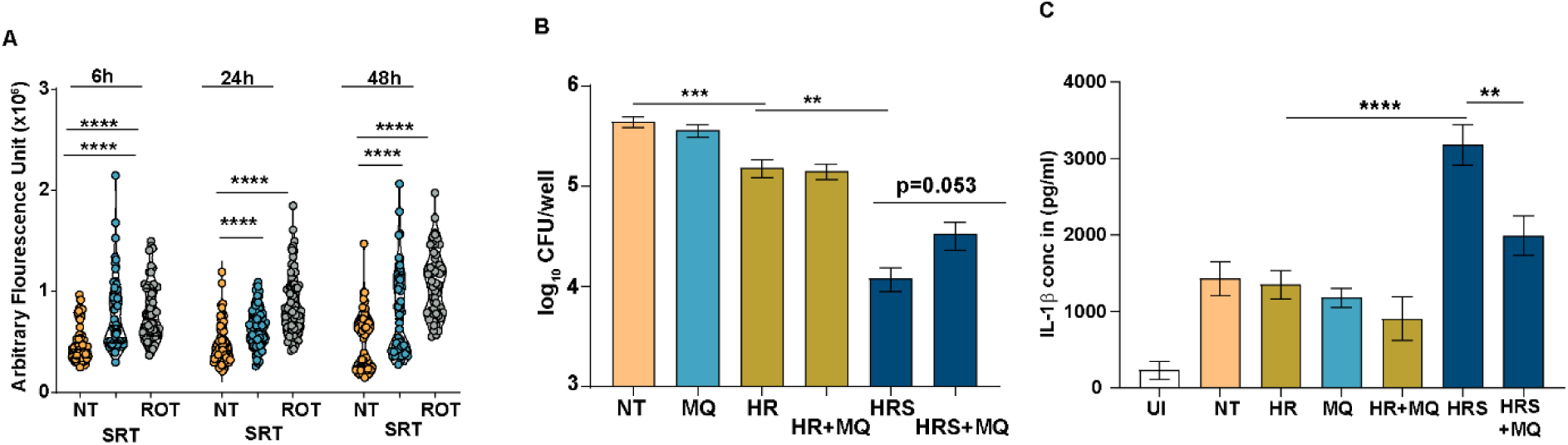
Mitochondrial ROS is important for efficient bacterial control in Mtb-infected macrophages. A). The levels of mitochondrial ROS induced in THP1 macrophages following treatment with SRT was estimated by staining with MitoSox. ROS values were quantitated as mean AFU±SEM of multiple cells/ image is depicted graphically from three independent experiments (N=3). B) The effect of inhibiting mitochondrial ROS by Mitoquinone (MQ) on bacterial control by Mtb-infected macrophages. Bacterial numbers in THP1 macrophages that were infected with Mtb and either left untreated (NT-B) or treated with HR, HR+ sertraline (HRS)at day0 and day3 post-treatment were enumerated by CFU plating and are represented as mean CFU ± SEM of triplicate assays from 3 independent experiments (N=3). C) The effect of addition of MQ on the levels of IL1β secretion in Mtb-infected macrophages was analyzed by ELISA and is represented as mean concentrations (pg/ml) ± SEM of triplicate assays from 3 independent experiments (N=3).

## Discussion

Therapies leveraging the host signalling pathways provide alternatives that circumvent the possibility of the development of drug-resistant bacterial strains in the population [3]. We have previously demonstrated the significance of adding the antidepressant sertraline to the standard TB therapy in achieving more efficient bacterial control and healthier tissue architecture. While we tested sertraline for its proven role in curbing the type I IFN response, we also demonstrated the importance of inflammasome activation in the enhanced bacterial control by sertraline [5]. By using sertraline to identify new mechanisms for increasing the potency of antibiotics on Mtb, we define the complete signalling cascade of inflammasome activation, a critical component of the innate immune system, to exert its antibacterial effects. Specifically, we reveal that the SRT mediated inflammasome activation culminates into an active K^+^ ion efflux, indispensable for the observed augmentation in antibiotic efficacy. In line with previous studies, our study also associates this outflow of K^+^ ion as a key regulator of type I IFN responses in macrophages [10]; inhibiting this efflux significantly enhances the infection-dependent type I IFN response significantly eroding the antimicrobial effect of the combination of sertraline and frontline TB drugs. These findings not only emphasize the significance of host inflammasome activation in Mtb control but also show its crucial role in antibiotic efficacy. Mtb infection has been associated with induction of IL1β via the NFκB pathway that has been implicated in improved Mtb killing directly as well as by activating alternate effector functions in murine and human macrophages [36, 37]. Activation of caspase-1 and caspase 4 plays a crucial role in inflammasome activation, and their inhibition leads to poor bacterial control [38]. Despite the combined inhibition of the two caspases, we did not observe a complete reversal in the HRS treated macrophages. It is plausible that the activation caspase-8 in Mtb infection could also mediate the GSDMD cleavage, potentially compensating for the loss of caspase-1 and caspase-4 activity [39].

Membrane pores created by GSDMD activation and oligomerization downstream of NLRP3 inflammasome activation function primarily in the release of mature IL1β from live innate cells [40–42]. Recently, GSDMD has been shown to have an affinity for cardiolipin that enables its binding to, and permeabilization of the bacterial cell membrane, leading to bacterial killing in the cytosol or within the phagosomes of phagocytes [43]. Additionally, GSDMD has been connected to targeting and eliminating cell-free bacteria when released from pyroptotic cells [44]. In fact, several intracellular bacterial infections, such as those caused by *Burkholderia thailandensis, Burkholderia cenocepacia, Francisella novicida,* and *Streptococcus zooepidemicus* initiate GSDMD-mediated pyroptosis leading to the elimination of pathogen by eradication of infected cells [45–47]. Although GSDMD can permeabilize and kill bacteria, chronic GSDMD activation results in excessive inflammation leading to tissue damage and cell death by pyroptosis [48].

A recent study has identified an important role for infection triggered GSDMD activation resulting in excessive necrosis and pyroptotic cell death as an important aspect of Mtb pathogenesis [49]. Our study highlights a novel mechanism induced by SRT that reorients phagocytic metabolism to an alternative non-lytic state by inducing significant proteolytic activation of GSDMD. In contrast to pyroptotic lysis, macrophages treated with SRT, favour the increased secretion of IL-1β observation and minimal necrosis as evidenced by the insignificant levels of LDH release [5]. Recent evidence suggests that GSDMD-activated macrophages can exist in two distinct states: a pyroptotic state characterized by cell lysis or a hyperactivated state with minimal cell lysis. In the hyperactivated state, increased levels of GSDMD membrane pores facilitate the release of mature IL-1β and other inflammatory mediators into the extracellular space. This is followed by membrane repair through the endosomal sorting complexes required for transport (ESCRT) pathway, which restores the damaged cell membrane[13]. This type of inflammasome mediated host defence has been demonstrated to facilitate control of another intracellular bacterial infection (*Salmonella typhimurium*) which allows the neutrophils to secrete IL-1β without compromising the cell membrane damage that induces pyroptosis [50].

We also define the ability of SRT to induce perturbation of mitochondrial physiology as a primary trigger for inflammasome activation in macrophages, establishing a critical link between mitochondrial function, inflammasome signalling and pathogen clearance. Mitochondria, as central hubs of cellular metabolism and immune signalling, play a crucial role in the observed effects of SRT and perturbation of physiology serves as a primary trigger for inflammasome activation, linking mitochondrial dynamics to immune regulation in TB. This finding is particularly significant given the emerging recognition of mitochondria as key regulators of innate immunity. While the reasons for SRT mediated mitochondrial dysfunction are still unclear and would be part of a comprehensive future study, our study confirms the amplification of the inflammasome response in enhancing host defence mechanisms against Mtb. As a result, the role of various gasdermin family members including GSDMD, which are considered key immune factors in defending against numerous infections, is now gaining increased attention. Our study thus underlines the importance of leveraging the synergistic effects of inflammasome activation and antibiotic treatment, for the development of more effective adjunct therapies for tuberculosis. Investigating other potential modulators of these pathways may reveal additional therapeutic targets for enhancing NLRP3 inflammasome activation and improving host defence mechanisms.

## Ethics

Human subjects: The study was conducted in strict accordance with recommendations of the National Ethical Guidelines for Biomedical and Health Research Involving Human Participants, Indian Council of Medical Research (ICMR), Government of India. The protocols followed were approved by the Institutional human ethics committee of IGIB, proposal no 10, 2016 and Ref no: CSIR-IGIB/IHEC/2017-18 Dt. 08.02.2018. Patient informed consent was obtained prior to commencing the work. Animal work the work was carried as per the requisites of the institutional animal ethics committee, approval (IGIB/IAEC/10/Nov/2023/05).

## Author contribution

AS, KB and VR were instrumental in the design of the work. AS and KB were involved in the conduct of the work. NY and RN were involved in the design and conduct of the ICPMS based detection of potassium ions in cells. None of the authors have any conflict of interest.

## Acknowledgements

The authors thank CSIR (VR-MLP2106, MLP2012) for supporting the study. CSIR-STS0016 is acknowledged for continuous maintenance of BSL3 and ABSL2 facilities. CSIR-BSC0403 is duly acknowledge for the microscopy facility. The student fellowships from CSIR-India (AS and KB) are acknowledged. Biorender.com is duly acknowledged for the graphical abstract preparation.

## References

1. Mase SR, Chorba T. Treatment of Drug-Resistant Tuberculosis. Clin Chest Med 2019; 40:775–95.

2. Nahid P, Mase SR, Migliori GB, et al. Treatment of Drug-Resistant Tuberculosis. An Official ATS/CDC/ERS/IDSA Clinical Practice Guideline. Am J Respir Crit Care Med 2019; 200:e93–e142.

3. Young C, Walzl G, Du Plessis N. Therapeutic host-directed strategies to improve outcome in tuberculosis. Mucosal Immunol 2020; 13:190–204.

4. Wong EB, Cohen KA, Bishai WR. Rising to the challenge: new therapies for tuberculosis. Trends Microbiol 2013; 21:493–501.

5. Shankaran D, Singh A, Dawa S, Arumugam P, Gandotra S, Rao V. The antidepressant sertraline provides a novel host directed therapy module for augmenting TB therapy. Elife 2023; 12.

6. Hwang HY, Shim JS, Kim D, Kwon HJ. Antidepressant drug sertraline modulates AMPK-MTOR signaling-mediated autophagy via targeting mitochondrial VDAC1 protein. Autophagy 2021; 17:2783–99.

7. Maes M, Song C, Lin AH, et al. Negative immunoregulatory effects of antidepressants: inhibition of interferon-gamma and stimulation of interleukin-10 secretion. Neuropsychopharmacology 1999; 20:370–9.

8. Szalach LP, Lisowska KA, Cubala WJ. The Influence of Antidepressants on the Immune System. Arch Immunol Ther Exp (Warsz) 2019; 67:143–51.

9. Chen S, Xuan J, Wan L, et al. Sertraline, an antidepressant, induces apoptosis in hepatic cells through the mitogen-activated protein kinase pathway. Toxicol Sci 2014; 137:404–15.

10. Mayer-Barber KD, Andrade BB, Barber DL, et al. Innate and adaptive interferons suppress IL-1alpha and IL-1beta production by distinct pulmonary myeloid subsets during Mycobacterium tuberculosis infection. Immunity 2011; 35:1023–34.

11. Bohrer AC, Tocheny C, Assmann M, Ganusov VV, Mayer-Barber KD. Cutting Edge: IL-1R1 Mediates Host Resistance to Mycobacterium tuberculosis by Trans-Protection of Infected Cells. J Immunol 2018; 201:1645–50.

12. Fremond CM, Togbe D, Doz E, et al. IL-1 receptor-mediated signal is an essential component of MyD88-dependent innate response to Mycobacterium tuberculosis infection. J Immunol 2007; 179:1178–89.

13. Evavold CL, Ruan J, Tan Y, Xia S, Wu H, Kagan JC. The Pore-Forming Protein Gasdermin D Regulates Interleukin-1 Secretion from Living Macrophages. Immunity 2018; 48:35–44 e6.

14. Zhou B, Abbott DW. Gasdermin E permits interleukin-1 beta release in distinct sublytic and pyroptotic phases. Cell Rep 2021; 35:108998.

15. Liu X, Zhang Z, Ruan J, et al. Inflammasome-activated gasdermin D causes pyroptosis by forming membrane pores. Nature 2016; 535:153–8.

16. Martinon F, Burns K, Tschopp J. The inflammasome: a molecular platform triggering activation of inflammatory caspases and processing of proIL-beta. Mol Cell 2002; 10:417–26.

17. Cruz CM, Rinna A, Forman HJ, Ventura AL, Persechini PM, Ojcius DM. ATP activates a reactive oxygen species-dependent oxidative stress response and secretion of proinflammatory cytokines in macrophages. J Biol Chem 2007; 282:2871–9.

18. Hornung V, Bauernfeind F, Halle A, et al. Silica crystals and aluminum salts activate the NALP3 inflammasome through phagosomal destabilization. Nat Immunol 2008; 9:847–56.

19. Shankaran D, Arumugam P, Vasanthakumar RP, et al. Modern Clinical Mycobacterium tuberculosis Strains Leverage Type I IFN Pathway for a Proinflammatory Response in the Host. J Immunol 2022; 209:1736–45.

20. Juffermans NP, Florquin S, Camoglio L, et al. Interleukin-1 signaling is essential for host defense during murine pulmonary tuberculosis. J Infect Dis 2000; 182:902–8.

21. McElvania Tekippe E, Allen IC, Hulseberg PD, et al. Granuloma formation and host defense in chronic Mycobacterium tuberculosis infection requires PYCARD/ASC but not NLRP3 or caspase-1. PLoS One 2010; 5:e12320.

22. Beckwith KS, Beckwith MS, Ullmann S, et al. Plasma membrane damage causes NLRP3 activation and pyroptosis during Mycobacterium tuberculosis infection. Nat Commun 2020; 11:2270.

23. Li H, Guan Y, Liang B, et al. Therapeutic potential of MCC950, a specific inhibitor of NLRP3 inflammasome. Eur J Pharmacol 2022; 928:175091.

24. Lopez-Castejon G, Brough D. Understanding the mechanism of IL-1beta secretion. Cytokine Growth Factor Rev 2011; 22:189–95.

25. Davis MA, Fairgrieve MR, Den Hartigh A, et al. Calpain drives pyroptotic vimentin cleavage, intermediate filament loss, and cell rupture that mediates immunostimulation. Proc Natl Acad Sci U S A 2019; 116:5061–70.

26. de Vasconcelos NM, Van Opdenbosch N, Van Gorp H, Parthoens E, Lamkanfi M. Single-cell analysis of pyroptosis dynamics reveals conserved GSDMD-mediated subcellular events that precede plasma membrane rupture. Cell Death Differ 2019; 26:146–61.

27. Zhu Q, Zheng M, Balakrishnan A, Karki R, Kanneganti TD. Gasdermin D Promotes AIM2 Inflammasome Activation and Is Required for Host Protection against Francisella novicida. J Immunol 2018; 201:3662–8.

28. Wang J, Deobald K, Re F. Gasdermin D Protects from Melioidosis through Pyroptosis and Direct Killing of Bacteria. J Immunol 2019; 202:3468–73.

29. Chai Q, Yu S, Zhong Y, et al. A bacterial phospholipid phosphatase inhibits host pyroptosis by hijacking ubiquitin. Science 2022; 378:eabq0132.

30. Banerjee I, Behl B, Mendonca M, et al. Gasdermin D Restrains Type I Interferon Response to Cytosolic DNA by Disrupting Ionic Homeostasis. Immunity 2018; 49:413–26 e5.

31. Won JH, Park S, Hong S, Son S, Yu JW. Rotenone-induced Impairment of Mitochondrial Electron Transport Chain Confers a Selective Priming Signal for NLRP3 Inflammasome Activation. J Biol Chem 2015; 290:27425–37.

32. Bronner DN, Abuaita BH, Chen X, et al. Endoplasmic Reticulum Stress Activates the Inflammasome via NLRP3- and Caspase-2-Driven Mitochondrial Damage. Immunity 2015; 43:451–62.

33. Neikirk K, Marshall AG, Kula B, Smith N, LeBlanc S, Hinton A, Jr. MitoTracker: A useful tool in need of better alternatives. Eur J Cell Biol 2023; 102:151371.

34. Marques E, Kramer R, Ryan DG. Multifaceted mitochondria in innate immunity. NPJ Metab Health Dis 2024; 2:6.

35. Shekhova E. Mitochondrial reactive oxygen species as major effectors of antimicrobial immunity. PLoS Pathog 2020; 16:e1008470.

36. Hackett EE, Charles-Messance H, O’Leary SM, et al. Mycobacterium tuberculosis Limits Host Glycolysis and IL-1beta by Restriction of PFK-M via MicroRNA-21. Cell Rep 2020; 30:124–36 e4.

37. Jayaraman P, Sada-Ovalle I, Nishimura T, et al. IL-1beta promotes antimicrobial immunity in macrophages by regulating TNFR signaling and caspase-3 activation. J Immunol 2013; 190:4196–204.

38. Zheng D, Liwinski T, Elinav E. Inflammasome activation and regulation: toward a better understanding of complex mechanisms. Cell Discov 2020; 6:36.

39. Schneider KS, Gross CJ, Dreier RF, et al. The Inflammasome Drives GSDMD-Independent Secondary Pyroptosis and IL-1 Release in the Absence of Caspase-1 Protease Activity. Cell Rep 2017; 21:3846–59.

40. Xia S, Zhang Z, Magupalli VG, et al. Gasdermin D pore structure reveals preferential release of mature interleukin-1. Nature 2021; 593:607–11.

41. Kuriakose T, Kanneganti TD. Gasdermin D Flashes an Exit Signal for IL-1. Immunity 2018; 48:1–3.

42. Tsuchiya K, Hosojima S, Hara H, et al. Gasdermin D mediates the maturation and release of IL-1alpha downstream of inflammasomes. Cell Rep 2021; 34:108887.

43. Miao R, Jiang C, Chang WY, et al. Gasdermin D permeabilization of mitochondrial inner and outer membranes accelerates and enhances pyroptosis. Immunity 2023; 56:2523–41 e8.

44. Jorgensen I, Rayamajhi M, Miao EA. Programmed cell death as a defence against infection. Nat Rev Immunol 2017; 17:151–64.

45. Aachoui Y, Kajiwara Y, Leaf IA, et al. Canonical Inflammasomes Drive IFN-gamma to Prime Caspase-11 in Defense against a Cytosol-Invasive Bacterium. Cell Host Microbe 2015; 18:320–32.

46. Estfanous S, Krause K, Anne MNK, et al. Author Correction: Gasdermin D restricts Burkholderia cenocepacia infection in vitro and in vivo. Sci Rep 2021; 11:12447.

47. Xu G, Guo Z, Liu Y, et al. Gasdermin D protects against Streptococcus equi subsp. zooepidemicus infection through macrophage pyroptosis. Front Immunol 2022; 13:1005925.

48. Vasudevan SO, Behl B, Rathinam VA. Pyroptosis-induced inflammation and tissue damage. Semin Immunol 2023; 69:101781.

49. Theobald SJ, Grab J, Fritsch M, et al. Gasdermin D mediates host cell death but not interleukin-1beta secretion in Mycobacterium tuberculosis-infected macrophages. Cell Death Discov 2021; 7:327.

50. Chen KW, Gross CJ, Sotomayor FV, et al. The neutrophil NLRC4 inflammasome selectively promotes IL-1beta maturation without pyroptosis during acute Salmonella challenge. Cell Rep 2014; 8:570–82.

